# Superabundant microRNAs are transcribed from human rDNA spacer promoters insulated by CTCF

**DOI:** 10.1101/2025.11.10.687711

**Authors:** Steven Henikoff, Jorja G. Henikoff

## Abstract

MicroRNAs are ∼22-nucleotide RNAs processed from primary transcripts and exported from the nucleus to repress gene expression by base-pairing to mRNAs. We find that the highest levels of RNA Polymerase II at human microRNA genes are within the ribosomal gene repeat arrays (rDNAs). Alignment of public nascent transcript data to the hs1 human genome assembly reveals a 50-nucleotide transcript for both miR-1275 and miR-6724, which exits from the nucleus with exceptional rapidity. We show that the miR-1275/miR-6724 transcription unit is closely flanked by CCCTC-binding factor (CTCF) within a <400-bp span of the rDNA spacer promoter. MiR-1275/miR-6724 and microRNA precursors expressed from the 5’ External Transcribed Spacer (5’ETS) are exported independently of known RNA processing activities and are detected in exosomes and as circulating cancer biomarkers. We propose that rDNA spacer promoter and 5’ETS microRNA genes have evolved for general regulatory functions in recipient cells.

## Introduction

MicroRNAs are small non-coding RNAs that regulate gene expression by base-pairing with mRNAs to prevent their translation or reduce their stability.^1^ Most microRNAs are processed from primary RNA Polymerase II (Pol II) transcripts as hairpin-like base-paired structures by the Microprocessor complex and exported from the nucleus, where the hairpin is cleaved by Dicer, and the processed microRNA is bound by an Argonaute protein. Some microRNAs are released from the cell within extracellular vesicles and taken up by recipient cells where they have been observed to alter gene expression.^2^

Many different microRNAs have been implicated in cancer, where they may regulate tumor suppressor or oncogene expression.^3-6^ The abundance of circulating microRNAs suggests that these are likely expressed at especially high levels and rapidly exported. However, the small size and rapid processing of microRNAs make them difficult to detect in clinical samples by standard RNA-sequencing (RNA-seq). Instead, microarray-based targeted assays are frequently used for cancer diagnostics^7,8^ as a substitute for deep sequencing of small RNAs.^9-11^ MicroRNA abundances can also be estimated using Cleavage Under Targeted Accessible Chromatin (CUTAC) with antibodies to Pol II in formalin-fixed paraffin-embedded (FFPE) sections.^12^ Using FFPE-CUTAC, we have shown that the conserved neural-specific miR-124a microRNA is abundantly expressed in normal mouse brain but strongly down-regulated in gliomas, a difference that was undetectable by RNA-seq. This finding encourages the expanded use of FFPE-CUTAC for microRNA detection in clinical applications, including cancer diagnosis, where a single assay will sensitively detect transcription of all RNAs, including mRNAs, microRNAs and other RNAs not easily detected in FFPEs.

Examination of our published human FFPE-CUTAC data revealed that a few microRNA loci consistently showed the highest RNAPII-Serine-5p (RNAPII-S5p) signals in all tissues and tumors tested when aligned to the hg19 or hg38 human genomic assemblies, where Serine-5 phosphorylation of the Rbp1 C-terminal domain marks Pol II that is paused before release for productive elongation.^13^ By aligning these microRNA genes to the full chm13 genome assembly we show that these and several other microRNA precursors are transcribed from the >200 ribosomal DNA repeats that reside on the short p arm of acrocentric chromosomes. Analysis of published run-on, nascent and small RNA sequencing data revealed that microRNA precursors are exported within minutes after release from RNA Polymerase and are not excised by the Microprocessor or cleaved by Dicer. Expression of a conserved 50-nucleotide hairpin transcript for two microRNAs is insulated by CCCTC-binding factor (CTCF) within the ribosomal DNA (rDNA) spacer promoter and exported with exceptional rapidity. Both miR-1275 and miR-6724 are abundant in exosomes and are the most differentially upregulated microRNAs in multiple human cancers. Our results suggest that microRNA precursors transcribed by Pol II from the rDNA spacer promoter and the 5’ External Transcribed Spacer (5’ETS) have evolved for gene regulatory functions within recipient cells.

## Results

### Paused RNA Pol II is superabundant at microRNA genes within human ribosomal DNA repeats

We previously applied RNAPII-S5p CUTAC to FFPEs from cancer patient samples representing seven diverse cancer types, and ranked candidate *cis-*regulatory elements (cCREs) by plotting the difference between tumor and normal samples as a function of the average on a log_10_ scale (modified Bland-Altman plots).^13^ Unexpectedly, three cCREs representing miR-663A were consistently the highest in overall abundance for all seven patients, showing little if any difference between tumor and normal samples (**Fig. 1A**). As miR-663A expresses a microRNA, we rank-ordered Pol II CUTAC signal over the 2,011 non-overlapping human microRNAs in RefSeq, which revealed that miR-663A ranks third behind miR-3648 and miR-10396 for all seven tumor and normal patient samples (**Fig. 1B**). We also ranked Pol II signal at microRNA sequences for the other FFPE-CUTAC samples from our previous study,^13^ plotting the average ranks for 15 breast and 30 meningioma patient samples, and again observed that miR-3648, miR-10396 and miR-663A were consistently the top three microRNA loci based on RNAPII-S5p abundance (**Fig. 1C-D**).

**Fig. 1:**
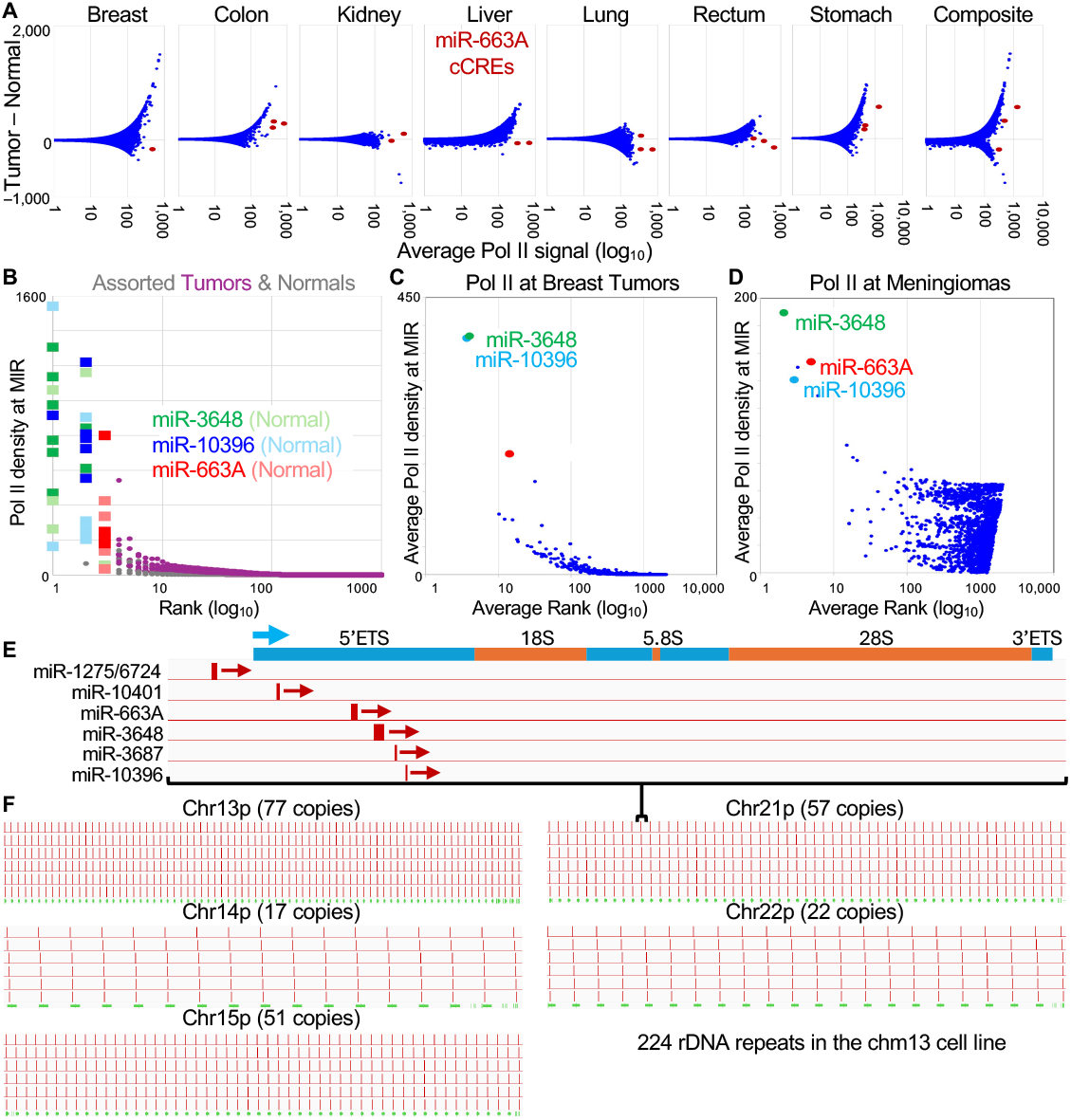
Pol II is superabundant at three microRNA genes in cancer and normal cells. (**A**) Modified Bland-Altman plots^7^ of Pol II FFPE-CUTAC data for 984,834 human cCREs from the ENCODE project^10^ from seven matched tumor and normal patient samples. MiR-663A is highlighted with up to three cCREs that show the highest average abundance but with no consistent differences between tumor and normal. (**B**) Pol II FFPE-CUTAC data were aligned to RefSeq-annotated microRNAs and rank-ordered, with miR-3648, miR-10396 and miR-663A ranking the highest with no consistent differences between tumor and normal. Ranks were averaged for patient samples from 15 breast tumors (**C**) and 30 meningiomas (**D**) and averages were plotted as a function of Pol II density, showing the same three microRNA genes consistently among the top three. Neither miR-3648 nor miR-10396 is represented by a cCRE, which accounts for their absence from the Bland-Altman plots in (A). (**E**) Alignment of miRBase-annotated microRNAs within the promoter and 5’ETS of the ribosomal genes for a representative repeat unit within Chromosome 21. (**F**) MegaBLAST^14^ was used to align microRNA sequences to hs1, which mapped them to each of the 224 repeat units in the hs1 genomic assembly.

RefSeq annotations for the hg38 genome assembly^14^ place miR-3648 and miR-10396 on Chromosome 21p, the short p arm of one of the five human acrocentric chromosomes, each of which harbors a tandem repeat array of ribosomal RNA (rRNA) genes. In 2021, the Telomere-to-Telomere (T2T) project released full genomic assemblies that resolved the tandemly repetitive satellite, telomeric and ribosomal gene arrays using long-read sequencing and improved genome assembly algorithms applied to the homozygous chm13 cell line.^15,16^ When we aligned the three microRNAs with superabundant RNAPII-S5p to the chm13 (hs1) genome assembly we found that all of them are located within the 5’ External Transcribed Spacer (5’ETS) of the ribosomal gene arrays (**Fig. 1E**). This implies that Pol II is not superabundant over these loci but rather is present at merely high abundance on a per locus basis, with the signal amplified by the 224 rDNA copies annotated in the chm13 cell line (**Fig. 1F**). miR-663A and miR-3648 were previously identified as residing within the 5’ETS by Yoshikawa and Fujii, who had searched annotated microRNAs^17^ against a single ribosomal DNA (rDNA) repeat unit.^18^ We confirmed their findings, adding miR-10396 and miR-10401 which were more recently annotated.^17^ In total, at least five microRNA genes reside in the 5’ETS, with miR-1275 and miR-6724 just upstream.

### A ribosomal DNA spacer promoter transcript rapidly exits from the nucleus

To obtain a profile of transcription that is not biased by RNA processing and does not depend on the particular RNA polymerase responsible, we aligned *in vitro* run-on transcription data to the hs1 genome assembly.^16^ We first analyzed data generated in various studies using Precision Run-On sequencing (PRO-seq), a run-on method that maps the RNA base in the active site of transcriptionally engaged RNA polymerases.^19^ By aligning PRO-seq data from the chm13 cell line to the hs1 assembly we observed a single sharply defined peak over each of the five acrocentric short arms, transcriptionally oriented 5’-to-3’ from telomere to centromere (**Fig. 2A**). Expansions of the scale for Chromosome 21 confirmed that the peak corresponded to the rDNA repeat array within Chr21p12 and that each rDNA repeat unit showed strong PRO-seq signals over the transcribed region (**Fig. 2C-E**). As ∼80% of transcription in the human genome is of the ribosomal genes,^20^ such extreme abundance of run-on transcription over the ribosomal RNA transcription unit was not unexpected. To confirm this *in vivo*, we turned to data from Transient Transcriptome sequencing (TT-seq) experiments, in which nascent RNA is pulse-labeled for ∼5 minutes with 4-thiouridine (4sU) followed by fragmentation, purification and sequencing of 4sU RNA.^21^ Using data from the DLD-1 colorectal adenocarcinoma cell line we observed the same five sharp peaks over the short p arms of the acrocentric chromosomes^22^ (**Fig. 2B**). These cells had been treated with 5,6-dichlorobenzimidazole (DRB), which inhibits the Cdk9 kinase of pTEFb and is important for productive elongation, consistent with enrichment of paused Pol II at microRNA genes within the rDNA repeats (**Fig. 2A-D**).

**Fig. 2:**
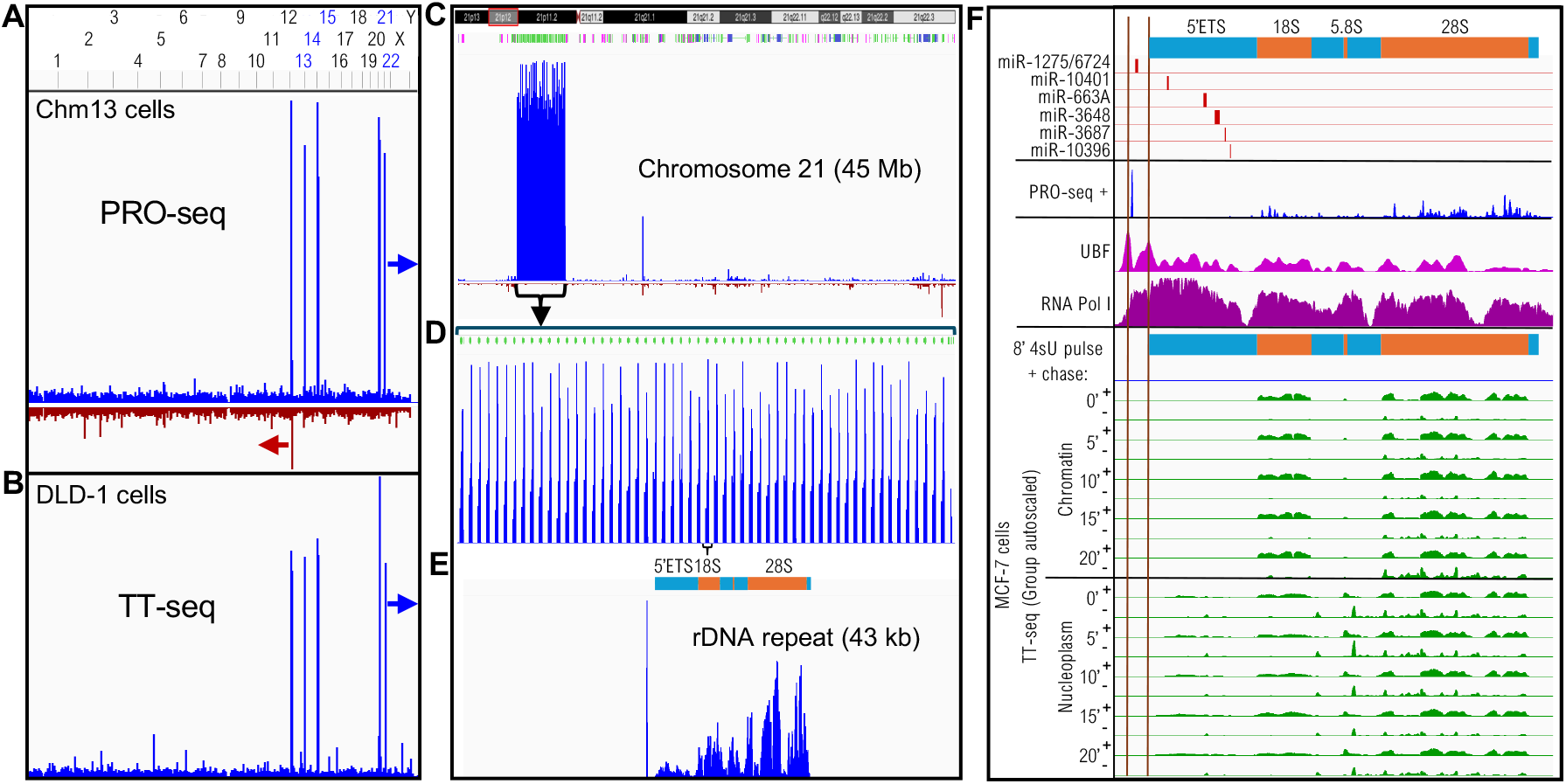
An rDNA spacer promoter transcript rapidly exits from the nucleus. (**A**) PRO-seq profile from the chm13 cell line shows strong peaks over the five human acrocentric chromosomes when aligned to hs1. The arrows indicate direction of transcription. (**B**) TT-seq data from DLD-1 cells confirms the PRO-seq forward strand profile *in vivo*. (**C**) The mini-chromosome from the hs1 assembly locates the strong PRO-seq peak over the hs1-annotated rDNA genes (green) within Chr21p12 (chr21:3,544,445-3,585,713). (**D**) Expansion of the Chr21 rDNA region. (**E**) Expansion of a single rDNA repeat unit centered over the transcribed region. (**F**) Top: Location of microRNA genes relative to the 47S rRNA precursor map within a representative rDNA repeat unit (chr21:3,544,445-3,585,713). Chm13 PRO-seq data^6^ (blue), UBF (magenta) and Pol I (purple) ChIP-seq data and chromatin-fractionated TT-seq pulse-chase data^8^ (green) below. Vertical lines indicate UBF promoter peaks that mark the spacer and 47S promoters.^12^

A conspicuous feature of the chm13 PRO-seq profile over the rRNA repeat unit is a sharp upstream peak that spans miR-1275 and miR-6724 (**Fig. 2E**). A similar pattern is observed over the rDNA repeat unit for PRO-seq data from Kasumi-1, a myeloblast cell line^23^ (**fig. S1 1-2**). The run-on and pulse-labeled transcripts over the ribosomal genes correspond to high levels of the Upstream Binding Factor (UBF) and Pol I (Ref. 24) (**fig. S1 3-4**) and relatively low levels of Pol II (**fig. S1 5-8**), as expected for transcription of the active rDNA genes by Pol I and repressive Pol II transcription of the inactive rDNA genes.^25^ The miR-1275/6724 PRO-seq peak lies within the UBF-defined rDNA spacer promoter.^26^ Total RNA-seq signal was high over 18S, 5.8S and 28S regions corresponding to stable transcripts^10^ (**fig. S1 9-10**), however DRB-resistant chromatin-enriched RNA^27^ was abundant primarily over the 5’ETS (**fig. S1 11-12**), consistent with rapid degradation of rRNA transcribed spacers.

To distinguish RNA polymerase release and export from programmed degradation, we examined TT-seq pulse-chase data from the MCF-7 breast cancer cell line, in which chromatin-bound and nucleoplasmic RNA fractions were sequenced separately.^25^ After an 8’ pulse of 4sU, the 18S, 5.8S and 28S regions show strong TT-seq signal in both fractions, and the 5’ETS region shows only nucleoplasmic RNA signal, however there is no TT-seq signal within the promoter and 5’ETS regions in the chromatin fraction (**Fig. 2F-G**). This suggests that the nascent run-on transcripts corresponding to rDNA promoter microRNAs were already exported from the nucleus before the end of the 8’ pulse. Transcripts on the opposite strand are seen throughout the 47S precursor region and may have resulted from low levels of Pol II transcription that have been observed to occur over silenced rDNA repeats.^24^ We infer that the abundant promoter microRNA transcripts observed in PRO-seq data are exported out of the nucleus within minutes, and the absence of TT-seq chromatin signal in the 5’ETS region suggests that these microRNAs are rapidly exported as well.

### The 50-nt rDNA spacer promoter microRNA precursor is present in other primates

To determine the precise length of the miR-1275/6724 nascent transcript, we compared the PRO-seq profile to the PRO-cap profile from the same DLD-1 cell line, where PRO-cap maps the modified 5’-end of the nascent transcript. We found that ∼95% of 5’-ends align precisely at the annotated 5’-end of miR-1275 and half of them have 3’ ends precisely at the 3’ end of miR-6724, consistent with the PRO-seq profile (**Fig. 3A**). The lack of excess nucleotides on either end implies that a 50-nucleotide microRNA precursor for miR-1275 and miR-6724 is a primary transcript that does not require cleavage activity of the Microprocessor complex.

**Fig. 3:**
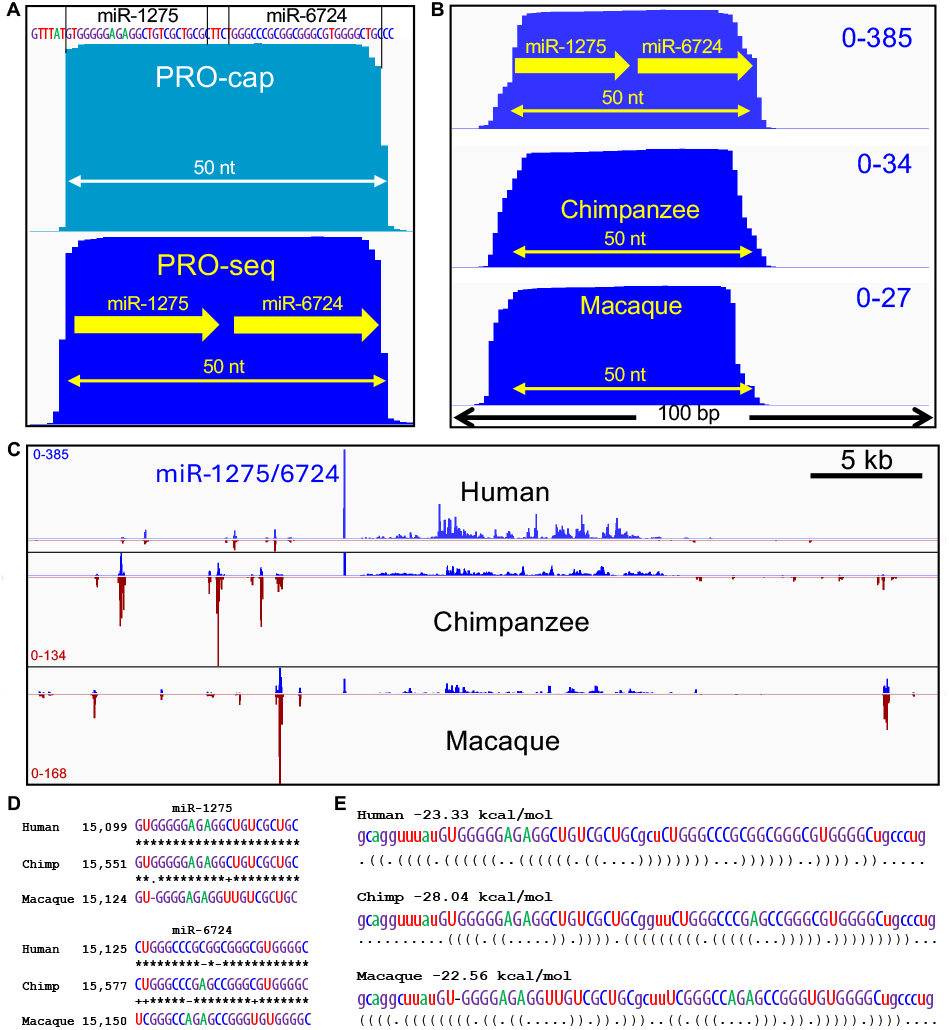
The 50-nt rDNA promoter microRNA precursor is found in other primates. (**A**) Forward-strand PRO-cap (turquoise) and PRO-seq (blue) mini-chromosome data for the rDNA spacer promoter transcription unit^2^. (**B**) A 100-bp region including the miR-1275/6724 peak at high resolution aligned by sequence similarity showing the + strand. (**C**) Forward-strand (blue) and reverse-strand (brown) PRO-seq data^4^ for a representative rDNA repeat unit (chr21:3,141,168-3,185,955) aligned with the corresponding regions of the chimpanzee rDNA repeat (GenBank KX061886.1) and the rhesus macaque rDNA repeat (KX061890.1) from CD4(+) T-cells. (**D**) Species alignments of the two microRNAs where each transition is indicated with a “+” and transversion or gap with a “–”. (**E**) The 50-bp human rDNA promoter hairpin structure is shown in Vienna format for each of the species.

A previous study identified miR-3648, miR-10396 and miR-6724 genes from hg38, where they had appeared to be uniquely present on Chromosome 21.^28^ The authors’ analysis of this region in ape and ancient human genomic assemblies at the time led them to suggest that this region of Chr21p11 emerged *de novo* in the human genome. Recently, T2T assemblies have been produced for several primates,^28^ which provide the opportunity to test whether the rDNA microRNAs have been maintained over evolutionary time. We aligned existing PRO-seq data from chimpanzee and rhesus macaque^29^ to their corresponding genome assemblies and observed that both species’ peaks precisely span the 50-nt miR-1275/6724 hairpin (**Fig. 3B-C**). There are few polymorphic differences between species (**Fig. 3D**), and the foldback structures that are substrates for cleavage and release to Argonaute proteins for translational regulation are also conserved (**Fig. 3E**), with similar minimum free energies of folding for all three species (∼20 kcal/mol)^30^ (**fig. S2**). We conclude that the microRNA cluster within the human ribosomal repeats has been functional at least since the last common ancestor of humans and macaques, ∼25 million years ago.

Evolutionary conservation of the rDNA promoter microRNA genes makes it most probable that they evolved to serve a housekeeping function requiring the high levels of transcript production that is possible with all >200 Pol I promoters producing the same transcript. For example, microRNAs found in extracellular vesicles have been shown to mediate changes in gene expression within recipient cells and are suspected in some cases to transmit drug resistance,^31^ where high abundance would give a microRNA-producing cell a competitive advantage.

### CTCF insulates the spacer promoter-miR-1275/6724 transcription unit

Zentner and co-workers^32^ mapped the CTCF transcription factor (TF) very close to the location of miR-1275/6724. To determine the precise location of this CTCF peak relative to miR-1275/6724, we analyzed DFF-ChIP data, in which native chromatin is fragmented using a non-specific double-strand endonuclease followed by immunoprecipitation.^33^ We separated fragments by size to distinguish CTCF that is directly bound to DNA (≤120 bp) from nucleosome-bound CTCF (>120 bp).^34^ We also plotted native DFF-ChIP data for TATA-binding protein (TBP),^35^ where previous work has shown TBP peaks over both the 47S rRNA promoter and the upstream spacer promoter corresponding to UBF peaks.^26^ We observed that directly bound CTCF maps both immediately upstream of TBP, and immediately downstream of the miR-1275/6724 PRO-seq signal (**Fig. 4A-B**). We also plotted native DFF-ChIP fragments for RNA polymerase-associated factors, Mediator, DRB Sensitivity Inducing Factor (DSIF) and Negative Elongation Factor (NELF), and observed precise co-localization with TBP over both spacer and 47S promoters, confirming that both promoters are active, although whether on the same or different rDNA copies could not be determined. The precisely defined squared-off DFF-seq peaks suggest that the promoter complexes (TBP-Mediator-DSIF-NELF) closely abut the RNA polymerase complex mapped by PRO-seq, and CTCF is bound immediately upstream of the promoter and downstream of the RNA polymerase, all within < 400 bp.

**Fig. 4:**
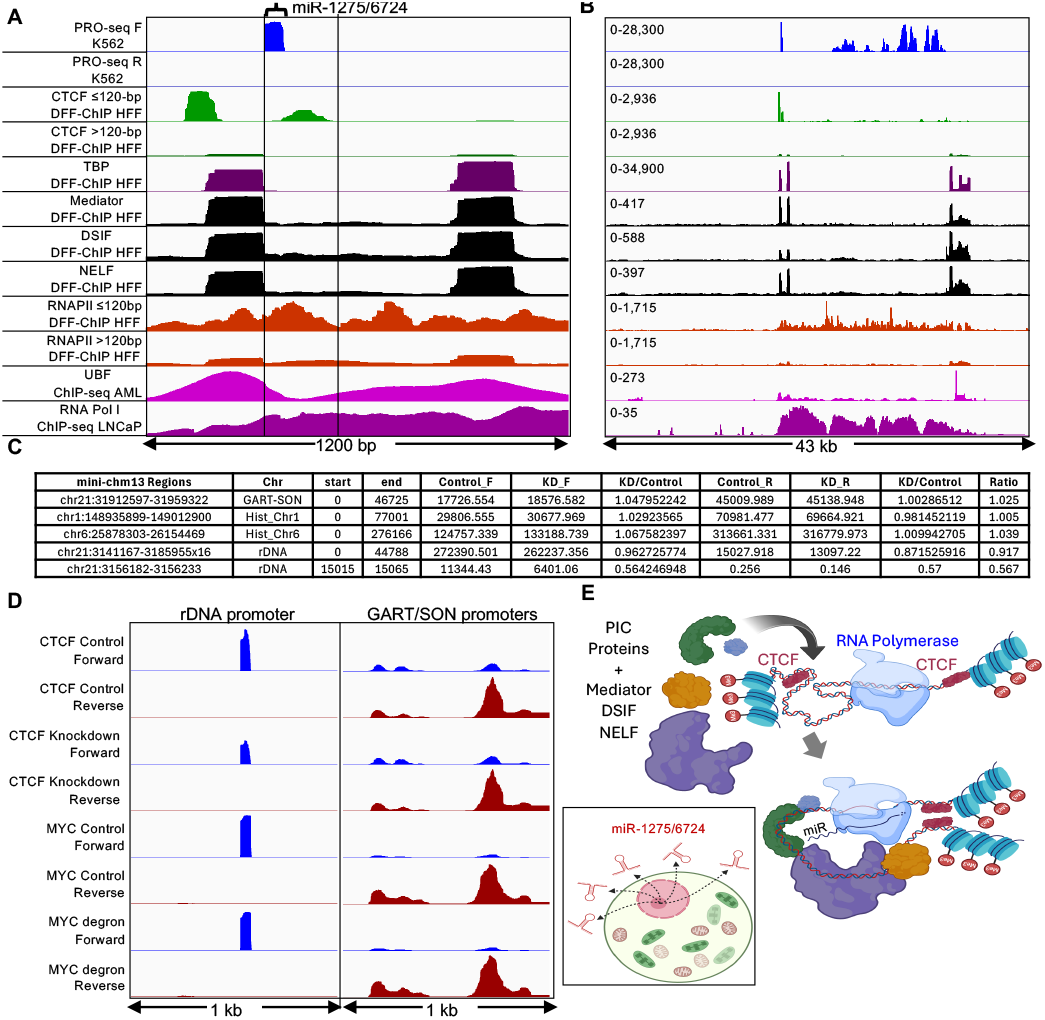
DFF-ChIP reveals precisely coordinated factors in the spacer promoter. (**A**) DFF-ChIP fragments from human foreskin fibroblast cells were aligned over the region spanning the rDNA spacer and 47S promoters, revealing the precise positioning of CTCF (green), TBP (purple), Mediator, DSIF and NELF (black) and Pol II (orange) relative to miR-1275/6724, UBF and RNA Pol I (vertical lines). (**B**) Same as (A) showing the full rDNA repeat unit and y-axis scales. (**C**) CTCF knockdown reduces total PRO-seq signal over miR-1275/6724 but has no effect on housekeeping control genes. Total PRO-seq signal for each strand was summed over entire mini-chromosomes and the fold-change was calculated for each strand and averaged for an overall ratio (last column). Controls were the GART-SON region (Ratio = 1.025), the histone gene cluster on Chr1 (Hist_Chr1, Ratio = 1.005) and the histone gene cluster on Chr6 (Hist_Chr6, Ratio = 1.039). A minor reduction with knockdown is seen for the entire rDNA repeat (Ratio = 0.917), and a substantial reduction for miR-1275/6724 (Ratio = 0.567). (**D**) Group-autoscaled PRO-seq profiles for 1-kb regions including the miR-1275/6724 promoter (left) and the bidirectional GART-SON promoter (right) ± depletion of CTCF or MYC. (**E**) CTCF insulates the <400-bp spacer promoter, for example from H3K27me3-marked nucleosomes, allowing for unimpeded Pre-Initiation Complex assembly and RNA polymerase engagement. The image was created with BioRender.com.

To further characterize the rDNA promoter region, we aligned Assay for Transposase-Accessible Chromatin with sequencing (ATAC-seq) data from Kasumi-1 cells and observed a peak just upstream of miR-1275/6728 (**fig. S3**). We also observed a peak of H3K27me3 just upstream of miR-1275/6724 with low-level H3K27ac in prostate cancer metastases, presumably from nucleosome-bound inactive copies. In addition, we aligned Cleavage Under Targets and Release Using Nuclease (CUT&RUN) data for the *MYC* oncogene, which is known to upregulate rDNA genes in mammals^36,37^ and Drosophila,^38^ and observed that in K562 cells, MYC and its partner MAX peak just upstream of miR-1275/6724, as did JUN in Hct116 cells (**fig. S3**).

To test whether CTCF is functionally important for miR-1275/6724 transcription, we examined CTCF knockdown data.^39^ This showed no effect on PRO-seq signal over housekeeping gene loci (knockdown/control = 1.01-1.03), a minor reduction over the rDNA repeat unit (0.92) and a substantial reduction over the 50-bp miR-1275/6724 locus (0.57), whereas degron-targeting of MYC showed little reduction of the miR-1275/6724 PRO-seq signal (**Fig. 4C-D**). On the basis of these observations from multiple cell lines, we infer that CTCF binding both immediately upstream of the spacer promoter and downstream of miR-1275/6724 insulates this intensely transcribed <400-bp region from nucleosome and/or UBF encroachment on either side (**Fig. 4E**). In support of this interpretation, the Cohesin complex was found to co-localize with the upstream peak,^40^ centered over the Zentner *et al*. CTCF motif,^32^ which is in the “forward” orientation for boundary pairing.^41^ The downstream peak spans an 83% GC-rich region with multiple regions potentially permissive for CTCF binding.^42^

### MiR-1275/6724 is regulated independently of rDNA

If CTCF functions to insulate miR-1275/6724, then it might also function to enforce independent regulation of the spacer promoter from the 47S rRNA promoter. To test this possibility, we first asked whether miR-1275/6724 uses a different RNA polymerase from Pol I, which transcribes the 47S rRNA genes. Alignment of Pol II high-resolution native DFF-seq fragments over the promoter region showed similar Pol II occupancy around both the spacer and 47S promoters extending through the entire transcribed rDNA, similar to what is observed for Pol I ChIP-seq (**Fig. 4A**). To test for possible Pol II transcription of miR-1275/6724, we asked whether Triptolide, which inhibits the catalytic activity of the Pol II-specific translocase complex, TFIIH, also reduces the miR-1275/6724 PRO-seq signal. We observed 1.4-to 2.1-fold increases depending on Triptolide concentration and treatment time for miR-1275/6724, with only slight changes up or down for 47S transcription in K562 cells (**Fig. 5A**), whereas control GART-SON housekeeping gene promoters showed drastic ∼30-fold reductions at the higher levels of inhibition (**Fig. 5B**).^43^ However, increased PRO-seq signal with Triptolide treatment was also seen in that study for certain long non-coding RNAs, consistent with classical *in vitro* reconstitution and recent cryoEM studies showing that TFIIH is not needed for promoter opening under torsional conditions favorable for spontaneous denaturation.^44,45^ This may explain the evident abutting of TBP-Mediator-DSIF-NELF on RNA polymerase at miR-1275/6724, where there is no free DNA available for action of the TFIIH translocase.

**Fig. 5:**
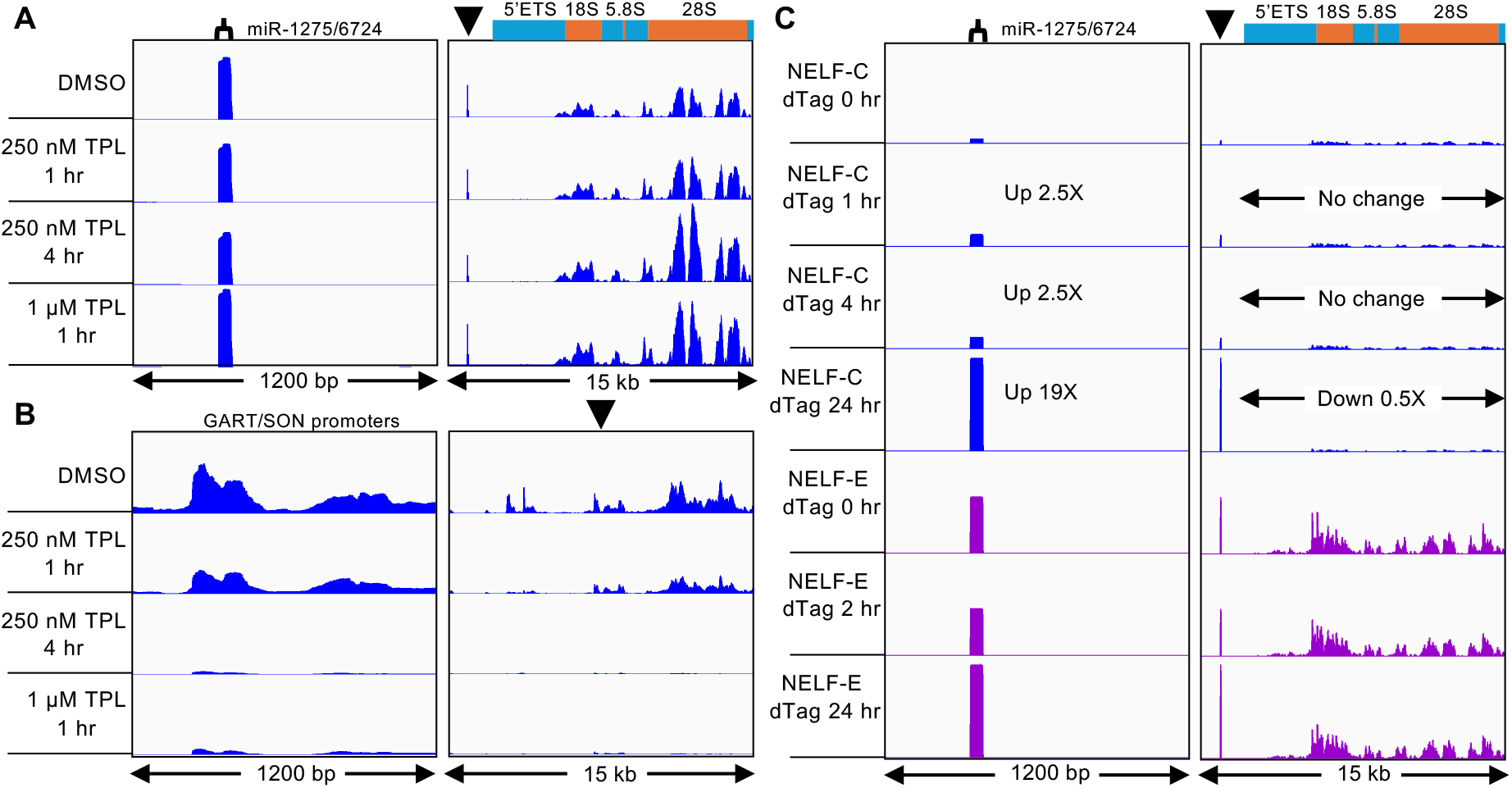
MiR-1275/6724 is regulated independently of the rDNA. PRO-seq data for TFIIH inhibition and NELF degradation. Group-autoscaled within series. (**A**) K562 cells^11^ were treated with Triptolide (TPL) at the indicated concentration and time, showing the rDNA promoter (left) and transcribed regions (right). (**B**) Same as (A) except for the control GART/SON promoter region (left) and zoomed out to 15 kb (right). DMSO, dimethylsulfoxide. (**C**) HFF cells had been transformed with NELF dTag constructs which were activated for degradation for the indicated times.^2^ NELF-C (blue); NELF-E (magenta).

Another Pol II-specific inhibitor is Flavopiridol, which like DRB, inhibits pTEFb, reducing productive elongation. When we analyzed PRO-seq data from human foreskin fibroblast (HFF) cells treated with Flavopiridol,^35^ we observed a 3.5-fold reduction at miR-1275/6724 (**fig. S4A**) and an ∼10-fold reduction of the GART-SON control promoters (**fig. S4C**). In contrast, the signal over the transcribed rDNA increased 1.5-fold (**fig. S4B**). The authors of this study also depleted TBP using a degron system over a 4-72 hour range,^35^ and we observed substantial increases in PRO-seq signal after Flavopiridol treatment for miR-1275/6724 and GART-SON but decreases for rDNA (**fig. S4**). Together, these observations imply that miR-1275/6724 is regulated independently of the ribosomal genes.

We next asked whether depletion of Negative Elongation Factor (NELF) would differentially affect miR-1275/6724 and rDNA transcription. On the basis of DLD-1 cell line data, we observed a 2.5-fold increase in miR-1275/6724 PRO-seq signal after a 1 hour dTag depletion of the NELF-C subunit, and 19-fold after 24 hours, by which time the rDNA PRO-seq signal had decreased by 50% (**Fig. 5C**). dTag depletion of NELF-E also increased the PRO-seq signal ∼2-fold without affecting the rDNA signal. These observations further demonstrate that miR-1275/6724 is regulated independently of the rRNA genes. The opposite effects on miR-1275/6724 and 47S transcription caused by inhibition of pTEFb and depletion of NELF, both of which are Pol II-specific initiation factors, would suggest that Pol II transcribes miR-1275/6724.

### microRNA precursors are expressed from the 5’ETS

The abundance of rRNAs in the 5’ETS undergoing degradation in the nucleoplasm may have obscured the production of microRNA precursors produced from this region in PRO-seq alignments (**Fig. 2F**). To detect these, we analyzed human Native Elongating Transcript Sequencing (NET-seq) data, based on in vivo nascent transcript 3’-end mapping of α-amanitin-treated cells following ribosomal RNA depletion.^46^ As α-amanitin is a specific inhibitor of the Rpb1 subunit of Pol II (Ref. 47) and has no effect on Pol I, (Ref. 48) NET-seq specifically reverse transcribes from 3’ ends of Pol II transcripts even when there is abundant Pol I transcription from other rDNA copies. For two promoter sites, including miR-1275/6724 and 10 sites within the 5’ETS, we observed ∼50-bp-wide broad peaks suggestive of microRNA precursors (**Fig. 6**). Broad ∼50-bp-wide peaks are seen to correspond in both HeLa S3 and HEK293T cells at 11 of the 12 sites. Some of these 5’ETS nascent transcripts are conspicuously abundant, including the broad peak spanning position 16,166-16,215 which suggests ∼200-fold higher abundance than miR-1275/6724 in both cell lines. As expected for interference from rRNA in PRO-seq data, we observed similar ∼50-bp broad peaks for the two spacer promoter sites in PRO-seq and PRO-cap data, but little if any resemblance between NET-seq and run-on data in any of the 5’ETS features.

**Fig. 6:**
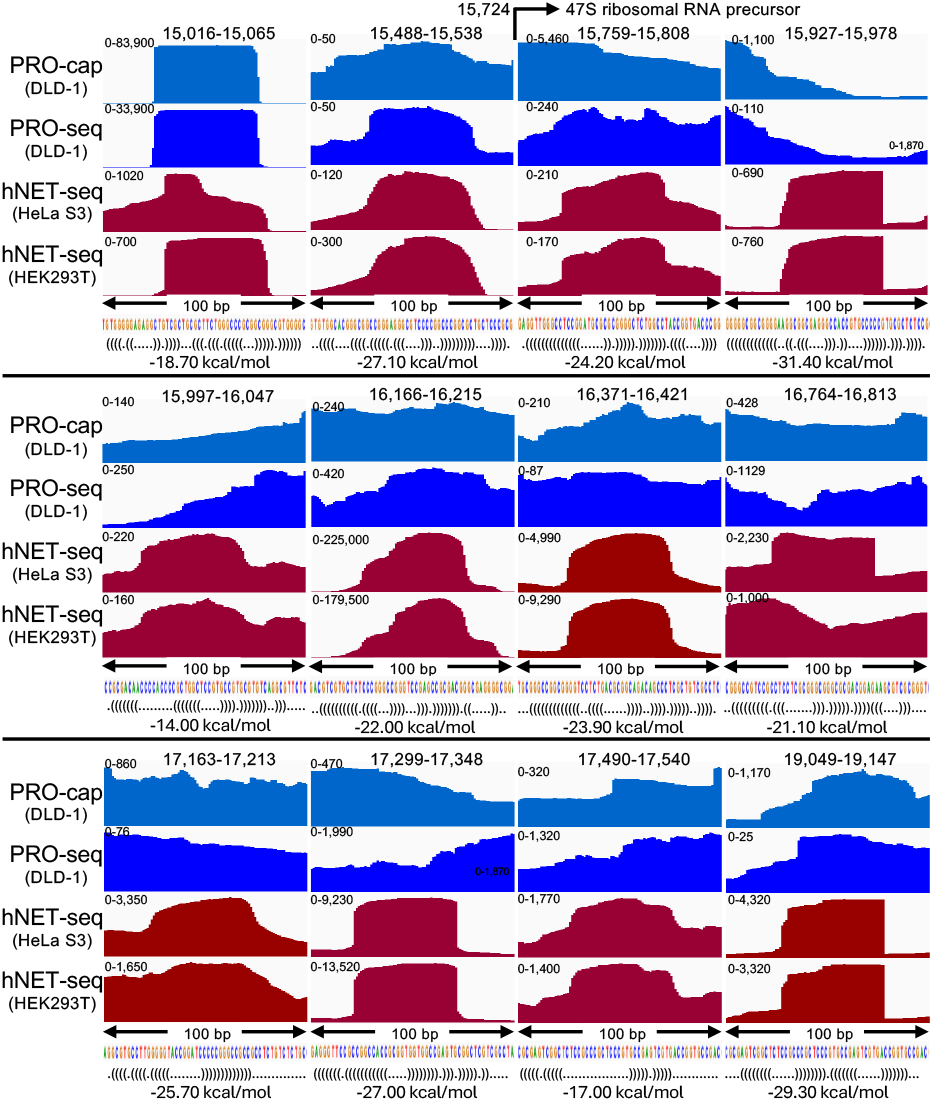
MicroRNA precursors from the 5’ETS are detected in a-amanitin-treated and rRNA depleted cells. Human NET-seq^13^ replicates from HeLa S3 and HEK293T cells were merged and aligned to the rDNA repeat unit. Each ∼50-bp wide peak feature was centered within a 100-bp window using Integrated Genome Viewer (IGV) for comparison to PRO-cap and PRO-seq data, autoscaled with the y-axis range shown at upper left. At top, each region is indicated by its position within the repeat unit, where 15,016-15,065 is the 50-bp span delimiting miR-1275/6724 and 17,163-17,213 includes miR-4466, whereas none of the other ∼50-bp peaks are annotated in miRBase. For each peak region, the sequence and predicted hairpin structure in Vienna format is shown, together with its RNAfold minimum free energy prediction.

To validate these ∼50-bp broad peaks as microRNA precursors, we used RNAfold,^30^ to identify potential hairpin structures. For all eleven putative microRNA precursors, RNAfold predicted hairpins comparable to that of miR-1275/6724, and for nine, the predicted minimum free energies of secondary structures were greater in magnitude than that of miR-1275/6724 (**Fig. 6**). Only one of these putative microRNA precursors, miR-4466, is annotated in miRBase, which is attributable to a matching G-rich 18-bp motif (GGGUGCGGGCCGGCGGGG) on Chromosome 6. This implies that the 5’ETS is a rich source of active microRNA precursors, overlooked because their gene sequences have been excluded from previous human genome assemblies, so that they can only be identified as microRNAs when total RNA-seq identifies a partial DNA sequence match in the small-RNA fraction.^49^

### rDNA chromatin and euchromatin levels are anti-correlated

Human rDNA copy numbers vary up to a few-fold between individuals (mean 469 ± 107),^50,51^ and we wondered whether rDNA copy number instability^20,52^ could be responsible for the discrepancies between levels of the rDNA promoter transcripts and both Pol I- and Pol II-dependent transcripts in response to different perturbations. To explore this possibility, we asked whether stable chromatin and transcriptional marks show concordance between the rDNA and other highly active loci in end-stage prostate cancer.^53^ In the case of H3K27me3, a mark of Polycomb silencing, we observed that their Cleavage Under Targets and Tagmentation (CUT&Tag) signals over rDNA and euchromatin on Chr1q transcribed by Pol II were almost perfectly anti-correlated (*r* = –0.999, *P* < 0.00001), whereas the histone gene clusters and the GART-SON housekeeping control regions were positively correlated with one another as expected (*r* = 0.377 to 0.732, **Fig. 7A, 7E**). Similarly, strong anti-correlations between rDNA and Chr1q euchromatin were seen for the active H3K27ac mark (*r* = –0.841, *P* < 0.00001, **Fig. 7B, 7F**) and for RNAPII-Ser5p signal from our FFPE-CUTAC data for multiple replicates from 13 breast tumors (*r* = –0.705, *P* = 0.0003, **Fig. 7C, 7G**) and 30 mostly benign meningiomas (*r* = –0.782, *P* < 0.00001, **Fig. 7D, 7H**). Such exceptionally strong anti-correlations for Pol I-transcribed rDNAs and Pol II-transcribed euchromatin for both active and silencing chromatin marks and paused Pol II are most easily understood if they reflect rDNA copy number differences between patients.

**Fig. 7:**
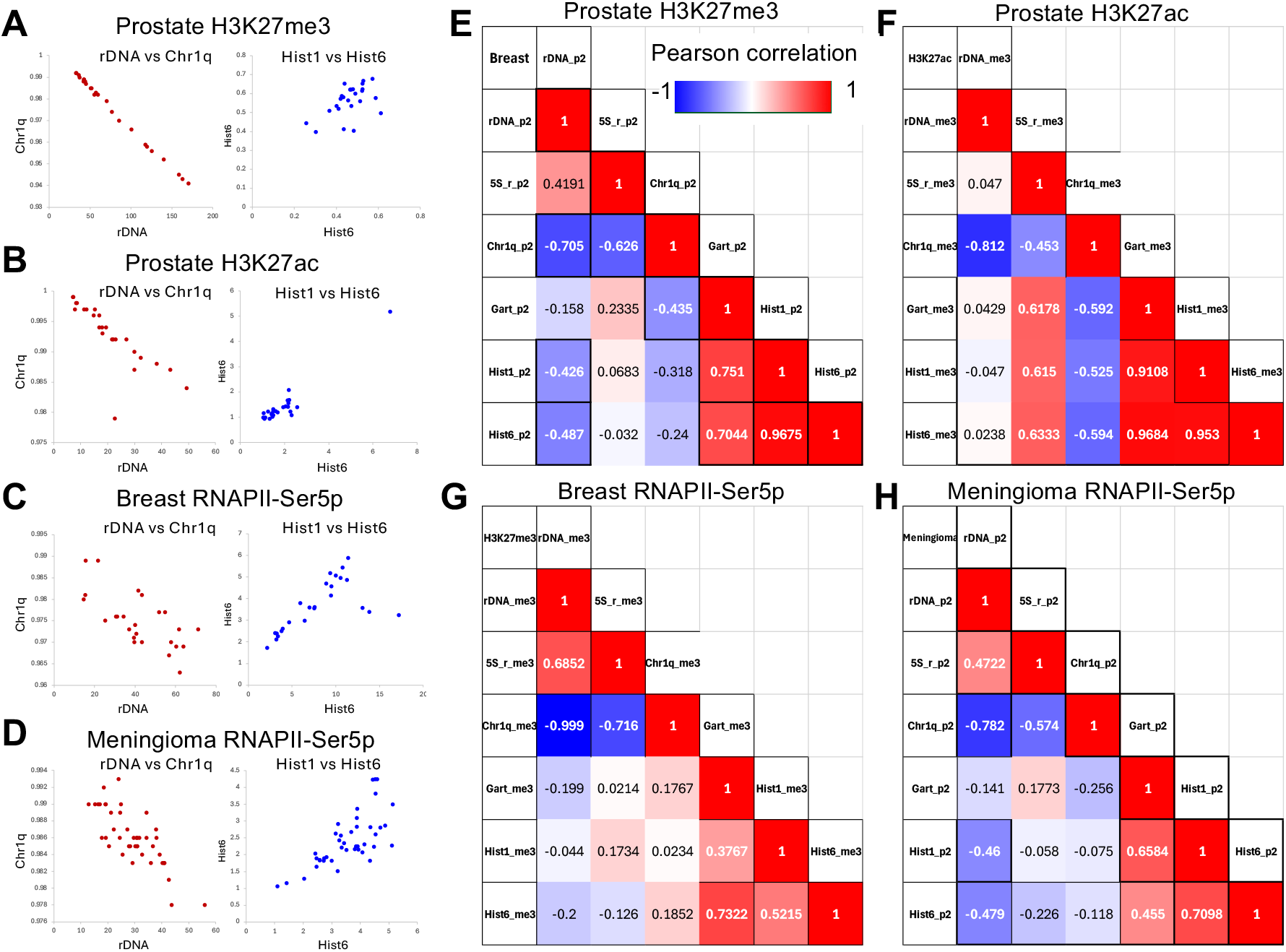
rDNA and euchromatin are anticorrelated for chromatin and Pol II marks. The human rDNA genes are almost perfectly anti-correlated with Chr1p euchromatin, from which the Hist1 cluster and the RNA5S cluster have been removed. (**A-B, E-F**) Correlation matrices for chromatin profiles of the silencing H3K27me3 mark (r = –0.999) and the active H3K27ac mark (r = –0.812) in multiple replicates from eight end-stage prostate cancer metastases. CUT&Tag patient data are from GSM8791938-GSM8791987 (Ref. 5) (**fig. S5**). Significant anti-correlations are also seen for multiple replicates of RNAPII-Ser5p in (**C, G**) 13 primary breast tumors (r = –0.705) and (**D, H**) 30 meningiomas (r - – 0.782) and between RNA5S and euchromatin (r = –0.453 to –0.782, p = 0.02 to 0.00006). In contrast, the Chr1 and Chr6 histone gene clusters are positively correlated with one another and with the control GART-SON locus in most other comparisons. FFPE-CUTAC data were from Ref. 9.

We also observed significant anti-correlations in patient data between the 5S rDNA genes, which are transcribed by Pol III, and Chr1q (r = –0.453 to –0.782, p = 0.02 to < 0.00001, **Fig. 7E-H**), although this cluster of 101 active Pol III transcribed genes reside on Chr1q. The significant positive correlation between rDNA and the 5S DNA for H3K27me3 (*r* = 0.69, *P* = 0.0002) and their anti-correlations with Chr1q euchromatin suggests that the two polymerases transcribing ribosomal RNA genes compete against Pol II for limiting nucleoside triphosphates.

### MiR-1275/6724 and 5’ETS microRNA transcripts use a non-canonical processing pathway

Most Pol II microRNA hairpin precursors are excised during transcription and are exported to the cytoplasm where the functional microRNA is released.^54^ However, a recent study reported that microRNA precursors processed by Pol III use a non-canonical pathway and are insensitive to inhibitors of the Microprocessor and Dicer.^55^ To determine whether rDNA promoter microRNAs are processed by the Microprocessor and Dicer, we aligned the small RNA-seq data from that study to the hs1 assembly and compared the profiles of rDNA microRNAs to that of miR-4521, which uses a non-canonical pathway. Inhibition of Pol III, which caused strong down-regulation of miR-4521, resulted in upregulation of the rDNA promoter microRNAs, miR-1275 and miR-6724 (**Fig. 8**). Upregulation with Pol III inhibition was also observed for the 5’ETS microRNAs, miR-10401 and miR-4466, which are predicted to form a tight hairpin structure (**fig. S5**). This opposite response to Pol III inhibition for rDNA microRNAs appears to be general, as two pairs of putative miRs in the 5’ETS, among those not annotated in miRBase (**Fig. 6**), also show upregulation with Pol III inhibition (**Fig. 8**). Altogether, six of the putative 5’ETS microRNA precursors identified in NET-seq data (**Fig. 6**) are confirmed to yield processed microRNAs that are upregulated with PolIII inhibition (**fig. S6**). Upregulation of miR-1275/6724 and 5’ETS microRNAs with reduction of Pol III is consistent with competition between RNA polymerases for limited nucleoside triphosphates as suggested above.

**Fig. 8:**
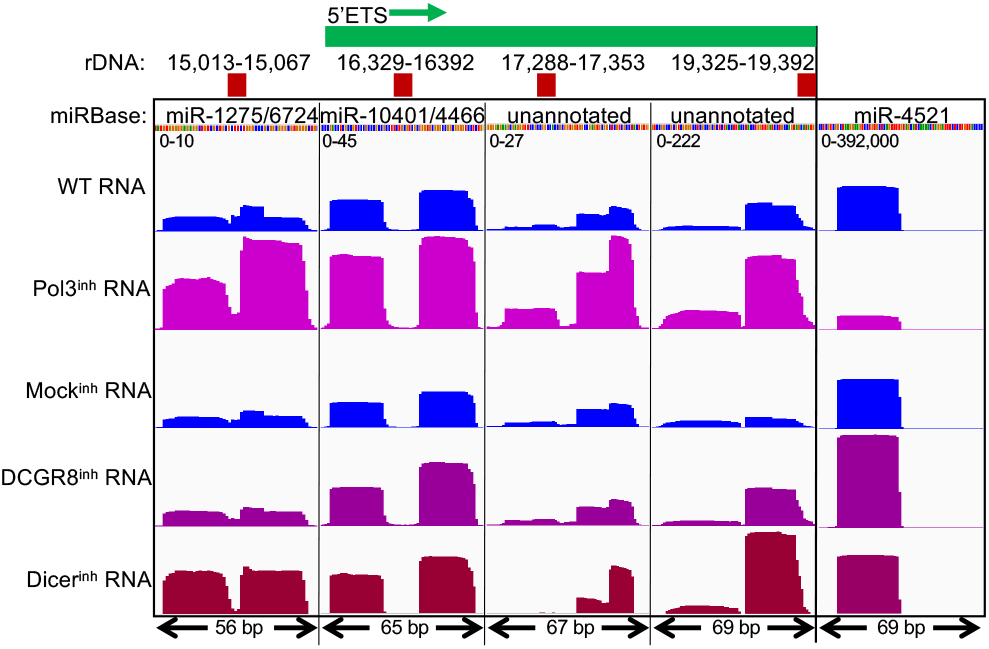
Processed microRNAs from the rDNA promoter and 5’ETS use a non-canonical processing pathway. Total small RNA-seq data from RKO colon carcinoma cells^1^ were aligned to a representative human rDNA repeat unit (chr21:3,141,168-3,185,955) and displayed in IGV format^3^, group-autoscaled for the five tracks, where the positions within the promoter-5’ETS region are indicated above. The scales range from 0 to 10 for miR-1275/6724 to 0 to 392,000 for miR-4521, consistent with rapid export of miR-1275/6724 from cells. Additional DCGR8- and Dicer-independent 5’ETS microRNAs are shown in fig. S6. Inhibition of the DCGR8 Microprocessor subunit and Dicer indicates that almost all of these microRNAs do not use the canonical processing pathways. WT, wild-type.

It is notable that the abundance of miR-1275/6724 is ∼1/40,000^th^ that of miR-4521, and 5’ ETS microRNAs are also orders of magnitude less abundant (**Fig. 8**). This is consistent with the amounts of rDNA-derived non-coding RNAs measured at <1/10,000^th^ the concentration of rRNA.^24^ We suggest that rapid export of primary transcripts from the cell and uptake by recipient cells can account for the low levels of processed rDNA spacer promoter and 5’ETS microRNAs relative to the level of miR-4521. Inhibition of DCGR8, a subunit of the Microprocessor complex,^54^ resulted in strong upregulation of miR-4521, and moderate upregulation of miR-1275, miR-6724, miR-10401 and miR-4466, although the effect on the unannotated transcripts varied from little or no effect to strong upregulation (**Fig. 8**). These observations exclude the possibility that the weak Pol III inhibitory activity of α-amanitin was responsible for generating 3’-hydroxyl ends in NET-seq experiments. The effect of inhibiting Dicer also resulted in upregulation of miR-4521 and mostly upregulation of the 5’ETS transcripts. Similarly, all six microRNA precursors from NET-seq data yielded microRNAs that are either increased in abundance or unchanged following either DCGR8 or Dicer inhibition (**fig. S6**). We conclude that the annotated microRNAs transcribed from the rDNA spacer promoter and 5’ETS are both Microprocessor- and Dicer-independent, using a non-canonical pathway for export and excision from the primary transcripts.

## Discussion

We have uncovered a unique mode of transcriptional regulation, in which the ribosomal gene spacer promoter is the source of a conserved 50-nt hairpin nascent transcript detected at exceptionally high levels in humans and other primates. Although our evidence is based on data that have been publicly available for years, we are not aware of any previous report of specific rDNA promoter transcripts in any organism. This oversight may be attributable in part to the fact that rRNA is by far the most serious contaminant of mRNAs in RNA sequencing studies, and standard protocols and analytical methods are designed to exclude it. Furthermore, the ribosomal genes have been deleted or masked in widely used eukaryotic genome assemblies and only with the hs1 genome assembly has it been possible to capture the full arrays on the short arms of all five human acrocentric chromosomes. Although the hs1 assembly has been available for ∼4 years, there are relatively few non-repetitive genes present in regions that are absent from previous genome assemblies, and so there has been little incentive to switch to it. We only noticed the superabundance of rDNA microRNAs as a puzzling anomaly in Bland-Altman plots, which required the full hs1 assembly to resolve. Overall, only 664 (1%) of Gene Expression Omnibus entries are aligned to hs1, compared to 26,133 for hg19 and 41,558 for hg38. Also, the small size and biased base composition of microRNAs result in their being attributed to single copy sequences that may occur by chance; for example, miR-1275 is annotated as a gene in Chromosome 6 euchromatin^17^ based on detection of a G-rich 17-bp motif (GUGGGGGAGAGGCUGUC) by small RNA-seq.^49^

Multiple lines of evidence suggest that the miR-1275/6724 rDNA spacer promoter transcript is rapidly exported from the nucleus: i) Despite showing the highest density of any nascent transcript in the rDNA using PRO-seq, transcription of the region is undetectable in both the chromatin-bound and the nucleoplasmic fractions after an 8 minute pulse of 4sU; ii) The two stable cleavage products of the 50-bp rDNA promoter transcript, miR-1275 and miR-6724, are detectable in cells at a level that is more than four orders of magnitude lower than that of miR-4521, and similarly low levels are seen for several 5’ETS microRNAs; iii) Both miR-1275 and miR-6724 have been recovered from exosomes and miR-1275 is abundant in human tissues, serum and urine.^7-9^

Many other microRNA transcripts are transcribed from within the 5’ETS region, including miR-10401, miR-4466, miR-663a, miR-3648, miR-3687 and miR-10396, as well as ten stable microRNA precursors not annotated in miRBase. All are transcribed from the same strand as the 5’ETS and are not distinguishable above background in PRO-seq data but are resolved in NET-seq data from α-amanitin-treated cells after rRNA depletion. Like rDNA spacer promoter microRNAs, 5’ETS microRNAs are found at levels that are orders of magnitude lower than that of miR-4521 and use a non-canonical processing pathway, suggesting that they are also rapidly exported, and many of them have been recovered in extracellular vesicles. Although these microRNA precursors are unique in being transcribed from the highly repetitive region rDNAs, evidently without being excised from a primary transcript, and are unusual in using a non-canonical processing pathway, they are structurally similar to ordinary microRNA precursors excised from primary Pol II transcripts. Therefore, we expect that their roles in gene regulation are no different from those of other exosome-enriched microRNAs, which are thought to regulate gene expression in recipient cells by base-pairing with cytoplasmic mRNAs in recipient cells.^1,56^

Both miR-1275 and miR-6724 have been detected at high levels in circulating microRNAs from patients with cancer and healthy individuals.^57,58^ In bladder cancer, circulating miR-6724 is the most significantly upregulated of the 2565 microRNAs assayed,^8^ and in hepatocellular carcinoma, circulating miR-6724 is the most significantly upregulated of the 2588 microRNAs assayed.^7^ MiR-1275 was found to be the most consistently upregulated of all microRNAs in tissue, plasma and urine samples, with a receiver operating characteristic curve area of 0.95 for urine, which confirms its exceptionally high abundance and diagnostic potential.^9^ Exosomes from prostate cancer cells containing miR-1275 have been shown to accelerate the proliferation of osteoblasts in culture.^6^ and upregulation of miR-1275 promoted the invasion ability of squamous cell carcinoma of head and neck, where elevated levels predict poorer patient outcome.^59^ Google Scholar lists 2,610 citations for “miR-1275 cancer”, and 1,780 citations for “mirR-1275 lung cancer”, with >1000 citations for exosomes and for five other common cancers, which suggests that the extraordinarily high output of this 50-nt primary transcript and of similar 5’ETS microRNA precursors is generally exploited by malignant cells. These observations support growing evidence that hypertranscription is a general cancer driver.^13,60^

## Materials and Methods

### Alignment and processing of sequencing datasets to the T2T-CHM13v2.0 (hs1) and chm13_mini reference sequences

#### Experimental Design

Datasets from public archives were downloaded and aligned to current genomic assemblies and derived mini-chromosomes to identify unannotated or mis-mapped microRNA precursors in chromatin-based and transcription-based epigenomic data.

#### Datasets and data processing

We downloaded fastq sequencing files from NCBI’s Short Read Archive (SRA) using their toolkit for every study analyzed. For single end sequencing, we then trimmed adapters using cutadapt 4.9 with parameters “-q20 -m20 -a AGATCGGAAGAGC” and aligned the trimmed reads to reference sequences using bowtie2 2.5 with parameters “--end-to-end --very-sensitive --no-unal --phred33”. For paired end sequencing we used cutadapt parameters “-m20 --nextseq-trim 20 -a AGATCGGAAGAGCACACGTCTGAACTCCAGTCA –A AGATCGGAAGAGCGTCGTGTAGGGAAAGAGTGT -Z” and bowtie2 parameters “--very-sensitive-local --soft-clipped-unmapped-tlen --no-unal --no-mixed --no-discordant --dovtail --phred33 -I 10 -X 1000”. Next, we extracted properly mapped reads from bowtie2 output into bed files of mapped fragments using the bedtools “bamtobed” command. When appropriate, we merged bed files for replicates using the UNIX sort command. Also, when appropriate we separated the bed files of mapped fragments into plus and minus strands, using the strand of READ1 for paired end reads. Finally, we computed fraction-normalized tracks from the final bed files of mapped reads using the bedtools “genomecov” command. Fraction-normalized tracks are the fraction of mapped counts at each base pair scaled by the size of the reference sequence so that if the counts were uniformly distributed there would be exactly one at each base pair. The UNIX scripts used are available at https://github.com/Henikoff/Scripts and https://zenodo.org/uploads/17737159.

### Alignment of sequencing datasets to the T2T-CHM13 (hs1) and chm13_mini reference sequences

Mini-chm13:

We constructed reference sequence chm13_mini from T2T-CHM13 version 2 (hs1) with these “chromosomes”:

GART-SON chr21:31,912,598-31,959,322 (46725 bp)

Hist_Chr1 chr1:148,935,900-149,012,900 (77001 bp)

Hist_Chr6 chr6:25,878,304-26,154,469 (276166 bp)

rDNA hs1 chr21:3,141,168-3,185,955 (44788 bp)

Illumina fastq sequencing files were downloaded from the NCBI SRA database. To cleanly separate euchromatin from both rDNAs on the acrocentric p arms and 5S rDNA on Chromosome 1q, for some analyses, we constructed reference sequence chm13_mini3 T2T-CHM13 version 2 (hs1) with these “chromosomes”:

GART-SON chr21:31912597-31959322

Hist_Chr1 chr1:148935899-149012900

Hist_Chr6 chr6:25878303-26154469

RNA5Sr chr1:227744287-228026142

rDNA chr21:3141167-3185955x16

Chr1q-remaining chr1:124048268-148935898 + chr1:149012901-227744286 + chr1:228026143-248387328

### Single-end sequencing

1. Trimmed adapters using cutadapt 4.9 with parameters: -q 20 -m 20 -a AGATCGGAAGAGC when appropriate
2. Aligned to reference sequences using bowtie2 2.5.4 with parameters: --end-to-end --very-sensitive --no-unal -q --phred33
3. Extracted mapped reads into bed file using bedtools bamtobed
4. Merged bed files for replicates (sort -m -k1,1 -k2n,2n -k3n,3n)
5. Separated bed files into plus/minus strands when appropriate (norm_strand.csh)
6. Computed fraction-normalized count tracks from bed files using bedtools genomecov (norm_beds.csh, fraction_norm.csh)

### Paired-end sequencing

1. Trimmed adapters using cutadapt 4.9 with parameters: -m 20 --nextseq-trim 20 -a AGATCGGAAGAGCACACGTCTGAACTCCAGTCA -A AGATCGGAAGAGCGTCGTGTAGGGAAAGAGTGT -Z
2. Aligned to reference sequences using bowtie2 2.5.4 with parameters: --very-sensitive-local --soft-clipped-unmapped-tlen --no-unal --no-mixed --no-discordant --dovetail --phred33 -I 10 -X 1000
3. Extracted mapped reads into bed file using bedtools bamtobed.
4. Merged bed files for replicates (sort -m -k1,1 -k2n,2n -k3n,3n)
5. Separated bed files into plus/minus strands based on the strand of READ1 when appropriate
6. Computed fraction-normalized count tracks from bed files using bedtools genomecov (norm_beds.csh, fraction_norm.csh)

## Datasets keyed to Figures

The following datasets were used in this study:

Fig. 1A-B : GSM8141208-GSM8141291

Fig. 1C : https://doi.org/10.5281/zenodo.13138686 DX_HS_P2S5_B*.bed.gz

Fig. 1D : https://doi.org/10.5281/zenodo.13138686 DX_HS_P2S5_M*.bed.gz

Fig. 2A-E , fig. S5 PRO-seq: SRX11346954-11346955

Fig. 2F : TT-seq: GSM6755736-6755745

Fig. 3A : GSM4296337, GSM4296339, GSM4296321

Fig. 3B-C Human: GSM2265095, GSM2265098, GSM2265100

Fig. 3B-C Chimp: GSM2265102, GSM2265104, GSM2265106

Fig. 3B-C Macaque: GSM2265108, GSM2265109, GSM2265111

Fig. 4A-B : GSM9050852-53, GSM8180606-7, GSM8839021-9022, GSM8180614-15, GSM8180610-11, GSM8180618-19, GSM8839019-9020, GSM9095398, GSM8267449

Fig. 4C-D : GSM6043500, GSM6043501, GSM6043506, GSM6043507

Fig. 5A-B : GSM9050852-53, GSM9050854-55, GSM9050856-57, GSM9050858-59

Fig. 5C : GSM4296314, GSM4296315, GSM4296316, GSM4296317, GSM4296318, GSM4296319, GSM4296320

Fig. 6 : GSM4296314, GSM4296337, GSM4296339, GSM1505438, GSM1505439, GSM1505440, GSM1505441

Fig. 7 : GSM8791938-8791987, GSM8141208-8141291

Fig. 8 WT: GSM9000286-9000288 (merged)

Fig. 8 Pol3-inh: GSM9000289-9000291 (merged)

Fig. 8 Mock-inh: GSM9000274-9000277 (merged)

Fig. 8 DCGR8-inh: GSM9000282-9000285 (merged)

Fig. 8 Dicer-inh: GSM9000278-9000281 (merged)

fig. S1–1-2 : GSM2552214-2552220 (merged)

fig. S1–3 : GSM9095398

fig. S1–4 : GSM8267447

fig. S1–5 : GSM8267449

fig. S1–6 : GSM8267450

Fig.S1–7 ,fig. S4 : https://doi.org/10.5281/zenodo.13138686 DX_HS_P2S5_BC1_0429.bed.gz

Fig.S1–8 : https://doi.org/10.5281/zenodo.13138686 DX_HS_P2S5_MA1_0401.bed.gz

fig. S1–9 : GSM8584120-8584125 (merged)

fig. S1–10 : GSM8584126-8584132 (merged)

fig. S1–11 : GSM1623148

fig. S1–12 : GSM1623146

fig. S2 : GSM3677824, GSM3677740-3677741, GSM36762-3677763, GSM3677784-3677785

fig. S3 : GSM5623305-5623317

fig. S5 : GSM8791938-8791987

## Supplementary Table 1

Mini-chm13

## Acknowledgments

We thank members of our lab, Toshio Tsukiyama and Steve Hahn for helpful discussions. This research was supported by the Howard Hughes Medical Institute (S.H.). Dedicated to the memory of Gabriel E. Zentner.

## Author contributions

S.H. and J.G.H. performed the analyses. S.H. wrote the original draft. S.H. and J.G.H. reviewed and edited the draft.

---

**fig. S1:**
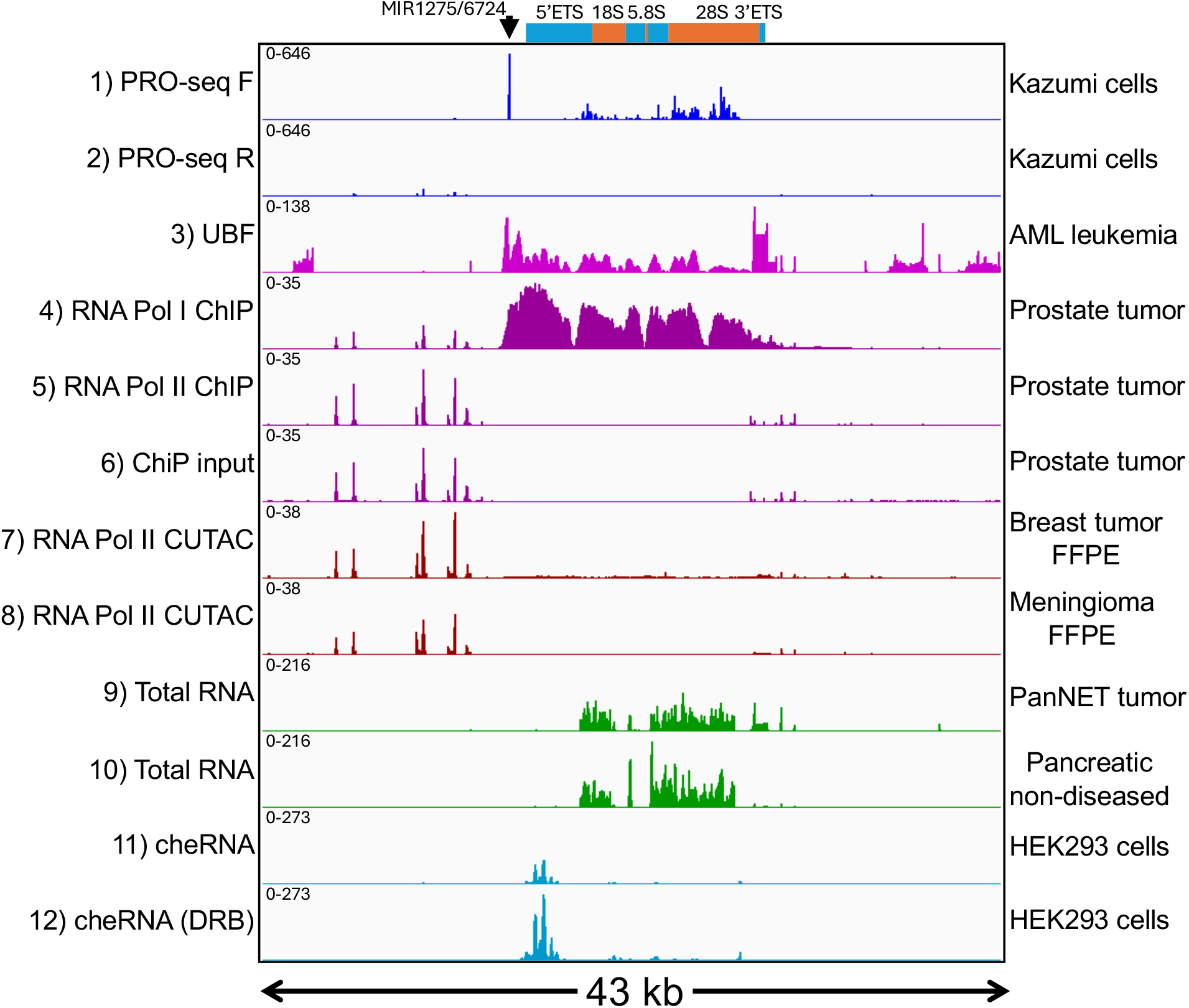
Multi-modal profile alignments over the human rDNA repeat unit. PRO-seq forward stand signal is high over the transcribed region with a peak of maximum height just upstream. (**1-2**) PRO-seq data from Ref. ^23^ within a representative rDNA repeat unit (chr21:3,544,445-3,585,713); (**3**) UBF ChIP-seq data from GSM9095398; (**4-6**) Pol I and Pol II ChIP-seq data from GSM8267449; (**7-8**) RNA Pol II FFPE-CUTAC data from Ref. ^13^; (**9-10**) Total RNA data from Ref. ^10^; (**11-12**) chromatin-enriched RNA data from Ref. ^27^.

**fig. S2:**
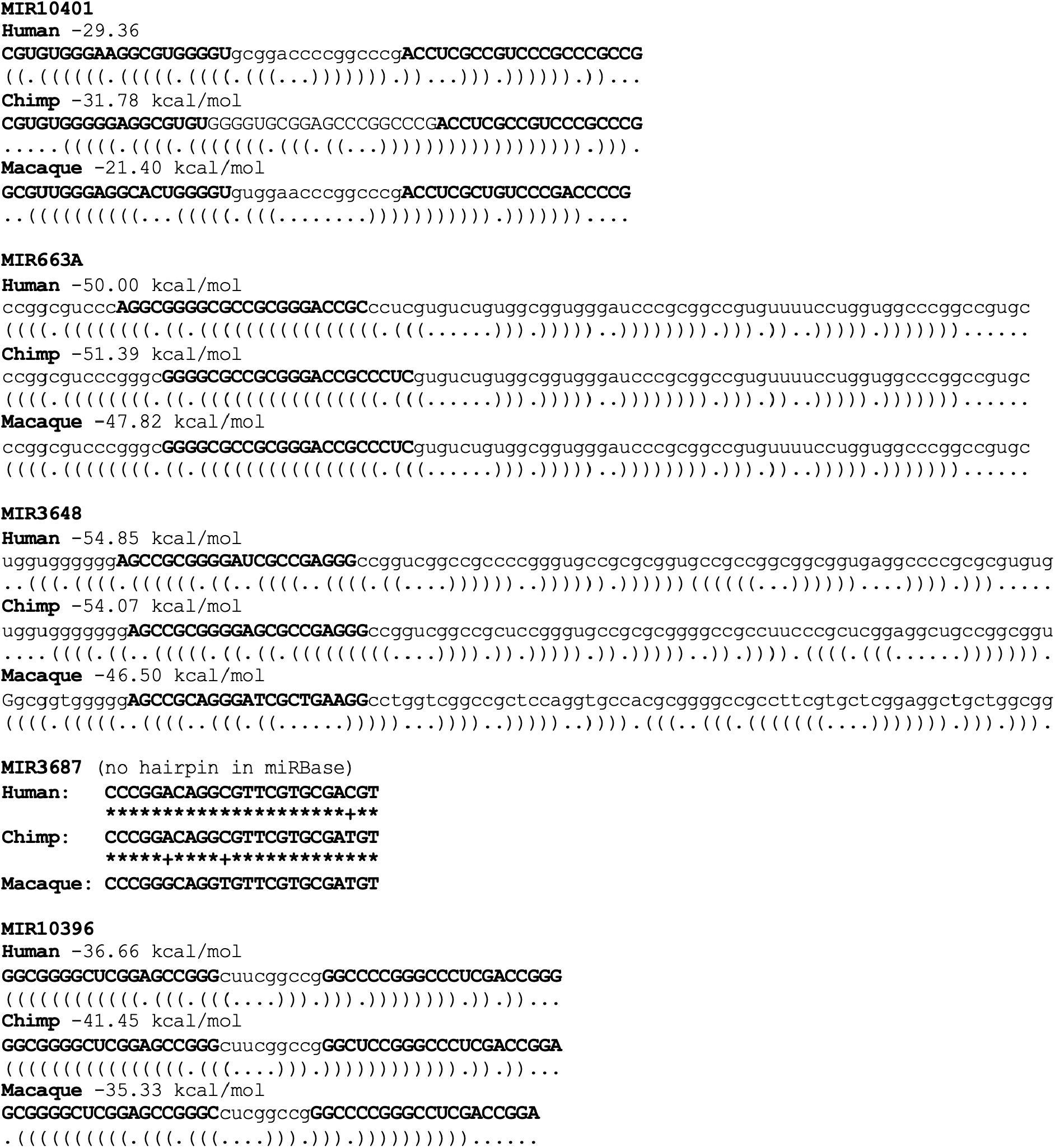
MicroRNAs in the 5’ETS region are conserved in primates. RNAfold^30^ was used to predict foldback structures for the five miRBase-annotated microRNAs in the 5’ETS region. Foldback structures are shown in Vienna format. expanded around the promoter and 5’ETS.

**fig. S3:**
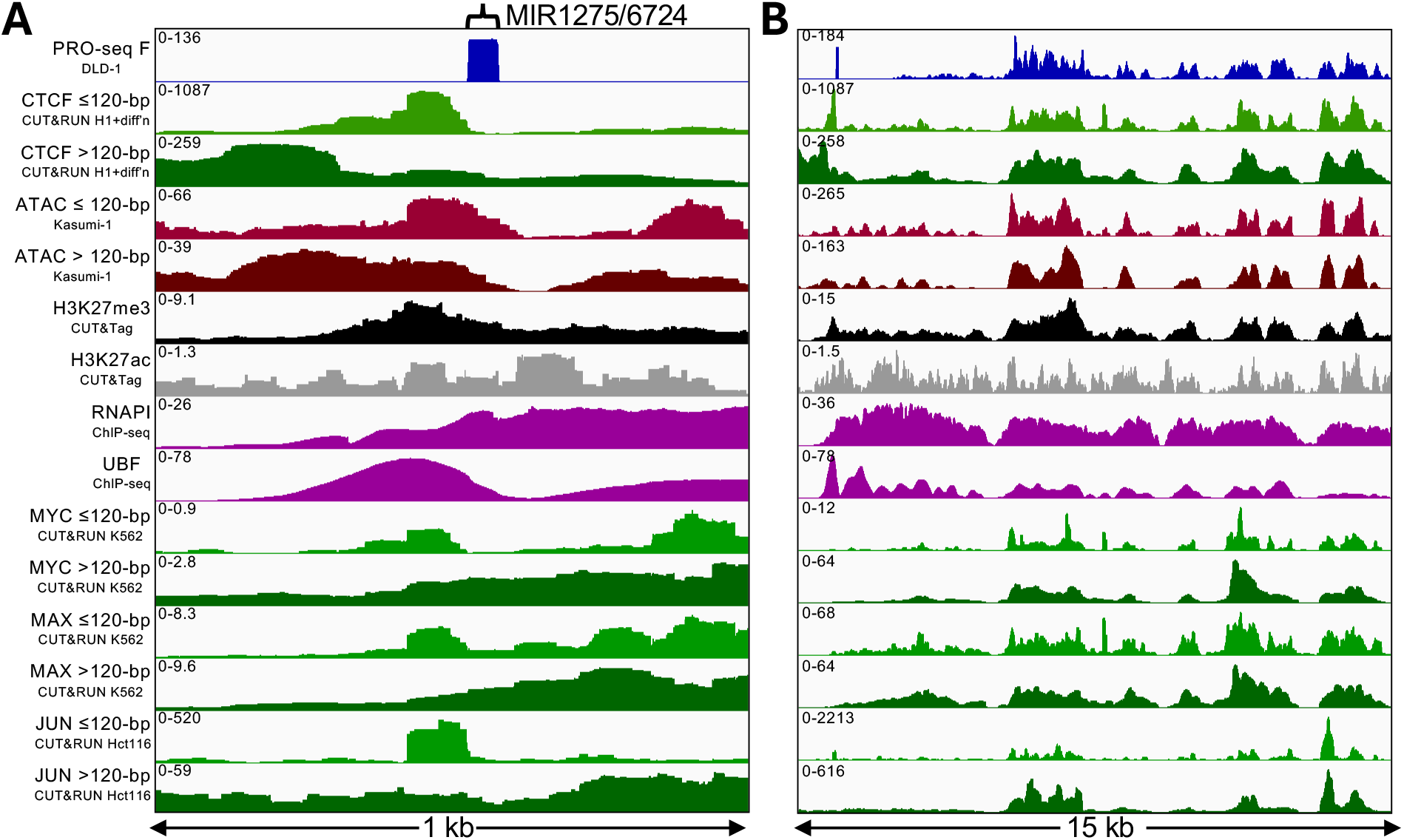
Myc/Max and AP-1 Transcription Factors bind to the miR-1275/6724 promoter. Subnucleosomal CTCF fragments (≤120 bp, green) map immediately adjacent to the PRO-seq peak (blue) directly over the miR-1275/6724 DNA sequence and the major UBF peak, whereas nucleosomal CTCF peaks (>120 bp, dark green) map farther upstream, with little overlap. Merged reads are from human H1 embryonic cells and 3-4 days of differentiation. A similar pattern is seen for ATAC-seq reads (light and dark brown). In metastatic prostate cancer, the mark of Polycomb silencing, H3K27me3 (black), but not H3K27ac (grey) a mark of active chromatin, peaks over the miR-1275/6724-adjacent CTCF and UBF sites, which suggests that the shift of CTCF from direct binding to nucleosome binding upstream is accompanied by gain of an H3K27me3 nucleosome that silences the rDNA spacer promoters. K562 cell MYC, MAX and Hct116 JUN CUT&RUN fragments (light and dark brown) were profiled as controls and show co-occupancy with UBF and shift downstream. (**B**) Same as (A) expanded around the promoter and 5’ETS regions.

**fig. S4:**
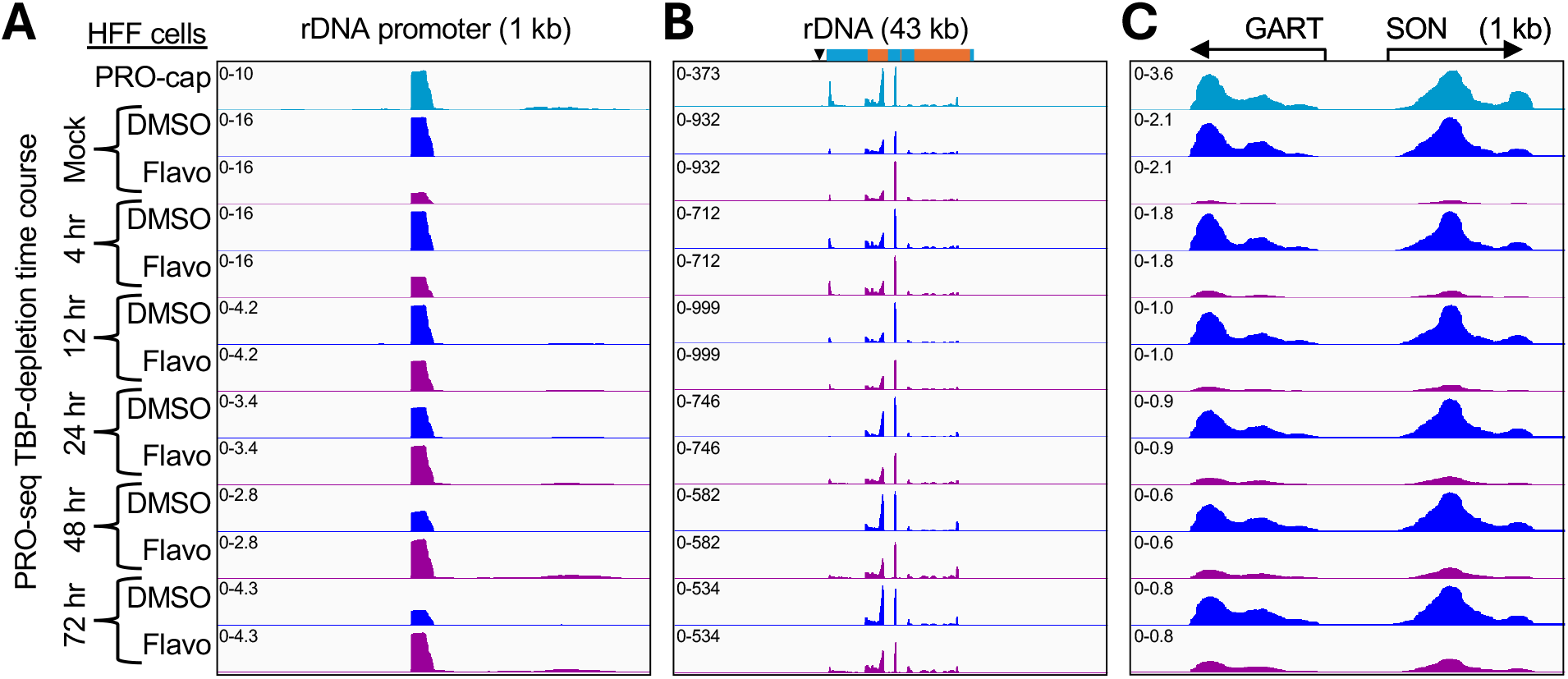
Forward-strand PRO-cap and PRO-seq mini-chromosome profiles at the rDNA promoter (**A**), the full rDNA repeat (**B**), and the GART-SON control region (**C**), for DMSO- or Flavopiridol-treated cultures after PROTAC induction for the indicated times.^61^ When TBP-depletion time points are autoscaled, dramatic PRO-seq signal increases after Flavopiridol treatment are seen for the miR-1275/6724, compared to modest decreases for the GART/SON bidirectional promoters and modest increases for rDNA transcription.

**fig. S5:**
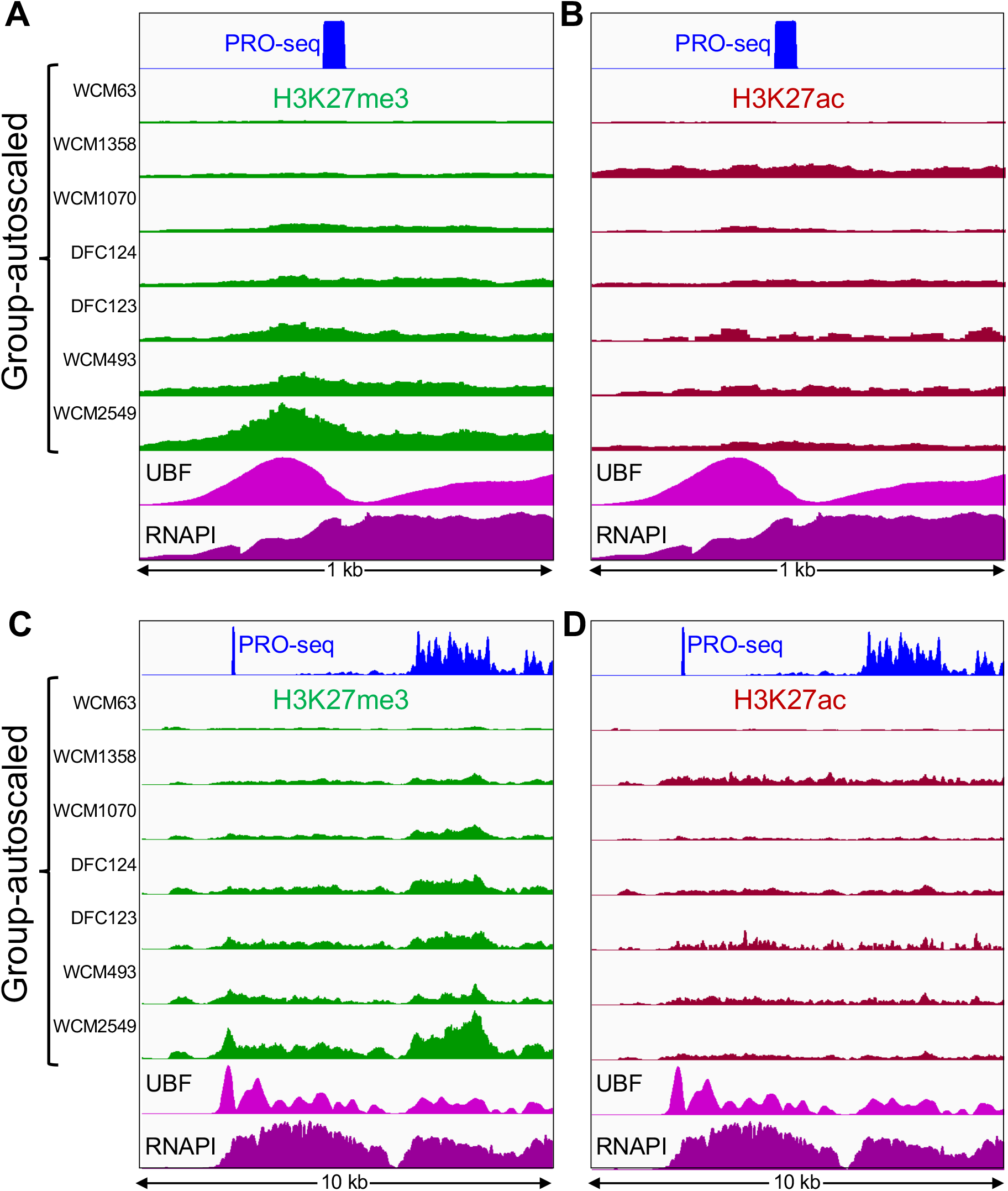
H3K27me3-marked rDNA genes vary in abundance between prostate cancer metastases. High H3K27me3 (**A**) and low H3K27ac (**B**) over the Pol I-transcribed regions from prostate metastases are consistent with densely packed Pol Is on active copies and heterochromatic histone marks on silenced copies. CUT&Tag patient data from GSM8791938-GSM8791987 (Ref. ^53^). Large variations in occupancy may reflect differences in both epigenetic and copy number differences between individuals.^50^ (**C-D**) Same as (A-B) expanded around the promoter and 5’ETS regions.

**fig. S6:**
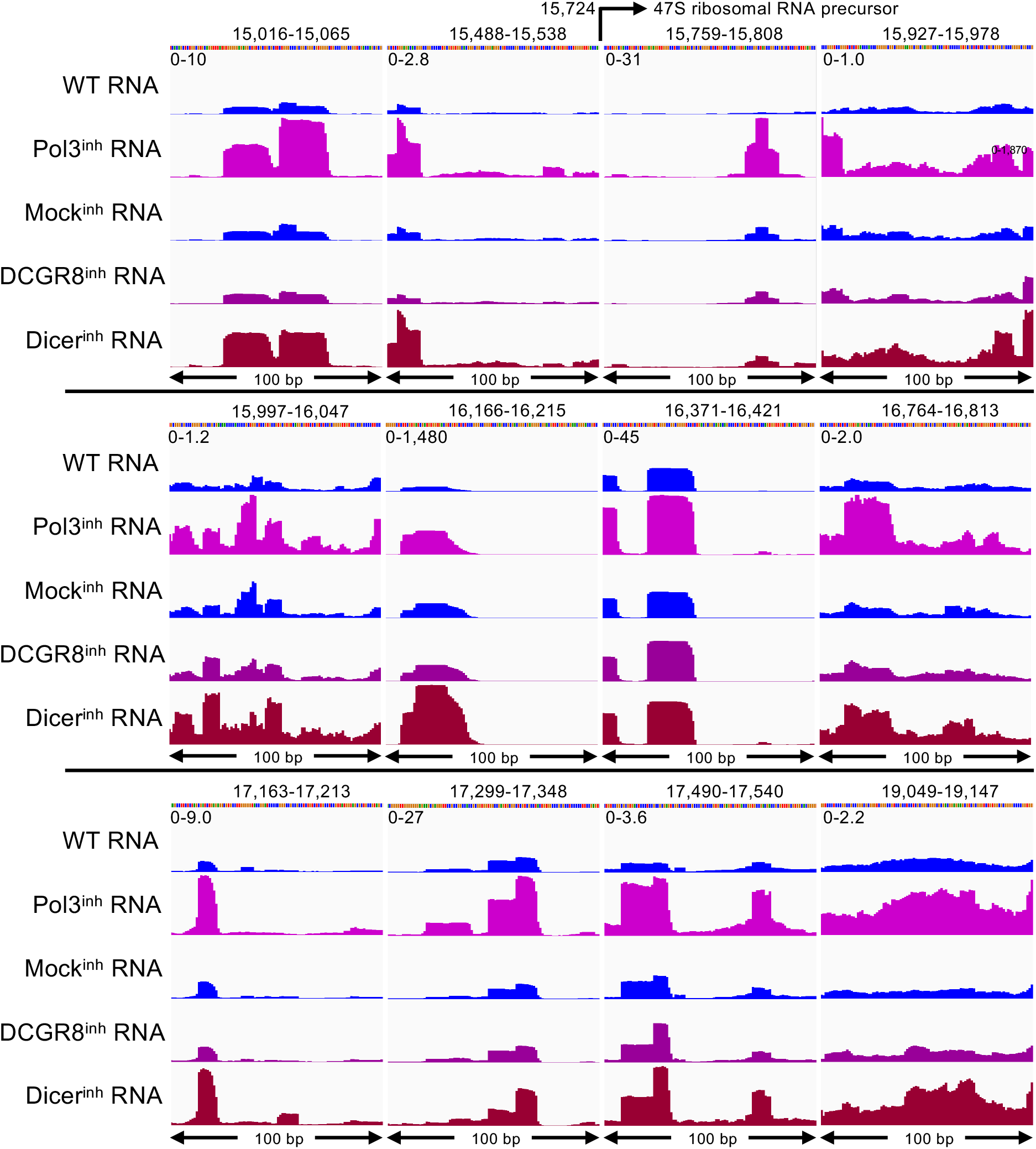
Processed microRNAs from the rDNA promoter and 5’ETS use a non-canonical processing pathway. See the legend to Fig. 8, using data from RKO cells.^55^ Group-autoscaled alignments were to the same 100-bp rDNA sub-regions used in the analysis shown in Fig. 6. Seven of the twelve microRNA precursors identified in NET-seq data show one or two characteristic fully processed microRNA peaks with signal counts ranging from 2.0 to 1,480, including miR-1275/6724, with 10 signal counts. Mir-4466 in region 16,371-16,421 is the only other annotated microRNA among those that show both NET-seq precursor signal and processed microRNA-seq signal. Inhibition of the DCGR8 Microprocessor subunit and the Dicer indicates that nearly all of these microRNAs do not use the canonical processing pathways.

